# Ultra-high field fMRI reveals functional patterns consistent with columnar organisation in human somatosensory cortex

**DOI:** 10.64898/2026.03.24.712494

**Authors:** Harriet Dempsey-Jones, Ashley York, Thomas B Shaw, Saskia Bollman, Markus Barth, Ross Cunnington, Alexander Puckett

## Abstract

In animal models, the primary somatosensory cortex (S1) exhibits columnar organisation, where vertically arranged neurons share functional properties. In humans, however, the thinness and folding of S1 have limited non-invasive investigations of such columnar structures. In this study, we aimed to identify columns in human S1 by delivering alternating bursts of 3 Hz and 30 Hz fingertip vibration while acquiring functional MRI time series at 7 Tesla. Using cortical surface modelling, we identified functional patterns in S1 that showed higher reliability, stronger differential responses, and greater statistical sensitivity than those observed in a frontal cortex control region (*p* = .001–.012 for reliability; *p* < .001 for differential signal; *p* = .004–.011 for sensitivity). Laminar analyses revealed depth-consistent frequency preferences in approximately 20–45% of S1 nodes, a pattern compatible with vertically organised functional structure. Although the relative signal difference between 3 Hz and 30 Hz was small (0.14% signal change), frequency tuning was reliably observed. Taken together, these findings reveal functional patterns in human S1 consistent with aspects of columnar-like organisation, providing non-invasive evidence of fine-scale functional architecture.

**Teaser:** fMRI reveals highly reliable but modestly selective responses in human S1, consistent with column-like functional organisation.

---

Over 60 years ago, Vernon Mountcastle’s pioneering single-cell recordings in cats revealed a striking “columnar” organisation in the somatosensory cortex (Mountcastle, 1957). By alternating stimulation of skin and deep tissues and recording from the first somatic region (the feline homologue of the primary somatosensory cortex), he observed that cells within a single electrode penetration, perpendicular to the cortical surface, responded preferentially to either form of stimulation. Across multiple, closely spaced sites, these responses appeared to form distinct columns of cells approximately 0.3–0.5 mm wide, sharing similar functional properties (Mountcastle, 1997). This work established a foundational architectural motif that has since been documented throughout the neocortex.

Subsequent animal studies identified columnar patterns in a wide range of cortical systems. In sensory cortex, ocular dominance and orientation columns are well characterised in vision (Hubel & Wiesel, 1959; reviewed in Yacoub et al., 2008). Columnar-like modules have also been described in frontal, limbic, and motor regions, where cortico-cortical projections terminate in vertically organised structures (Goldman & Nauta, 1977). Their prevalence has led to the proposal that columnar organisation may reflect a general organising principle of the neocortex (Mountcastle, 1997). These findings naturally motivate attempts to examine analogous fine-scale organisation in the human brain.

Recent advances in high-resolution neuroimaging have made it increasingly feasible to study fine-scale cortical organisation in humans. Early work at high magnetic field (4 Tesla) demonstrated that functional MRI could resolve submillimetre-scale spatial patterns (Menon et al., 1997), and subsequent developments at ultra-high field (7 Tesla) have substantially improved both resolution and sensitivity (Nasr et al., 2016; Yacoub et al., 2008). These advances have enabled the non-invasive identification of columnar-scale patterns such as ocular dominance and orientation tuning in human visual cortex (Cheng et al., 2001; Menon et al., 1997; Yacoub et al., 2007, 2008).

Despite these successes, investigating columnar organisation in human primary somatosensory cortex (S1) remains challenging. Compared to visual cortex, S1 is more highly folded and relatively thin (Fischl & Dale, 2000), complicating analyses across cortical depth. The delivery of precise, controlled tactile stimulation in the MR environment also requires specialised MR-compatible devices with careful validation (Puckett & Sanchez Panchuelo, 2023; Travassos et al., 2023). Although S1 was the first area in which cortical columns were discovered, establishing comparable non-invasive evidence in humans remains challenging.

Physiological studies in non-human primates provide important guidance for identifying the types of functional distinctions that might reveal columnar organisation. Early work reported “rapidly adapting” S1 cortical columns that responded at the onset and offset of 1-second skin indentations, interleaved with “slowly adapting” columns that responded throughout these periods of indentation (Sur et al., 1981, 1984); see also (Dykes et al., 1980). While slowly adapting columns appeared to span only the middle layers of cortex, rapidly adapting columns extend over all layers (Sur et al., 1984). Later work identified slowly adapting, rapidly adapting, and rapidly adapting type II (Pacinian) columns responding to different speeds of vibration stimulation (1 Hz, 30 Hz and 200Hz, respectively) (Chen et al., 2001; Friedman et al., 2004). However, it has also been proposed that such distinctions may reflect perceptual categories (e.g., “tap,” “flutter,” “vibration”) rather than strict peripheral receptor classes (Saal & Bensmaia, 2014). Regardless of the underlying organisational mechanism, applying vibrational stimuli at distinct frequencies provides a principled approach to probe potential columnar patterns in human S1.

Here, we used ultra-high field (7T) MRI and cortical surface modelling to investigate functional patterns in S1 during vibrotactile stimulation of the fingertips at 3 Hz and 30 Hz – frequencies chosen to preferentially engage slowly and rapidly adapting pathways and to elicit distinct perceptual sensations (tap and flutter). We first generated cortical maps reflecting frequency preference, averaged across cortical depths, and assessed their reproducibility across independent scanning blocks (Part 1). We then examined a defining characteristic of columnar organisation: consistency of functional preference across cortical depth (Part 2). Specifically, laminar analyses tested whether frequency preferences observed at the pial surface were preserved down to the grey–white boundary. Together, these analyses provide a non-invasive test of whether functional patterns in human S1 are consistent with aspects of columnar architecture described in animal models.

## Results

Participants performed two tasks involving vibrotactile stimulation to the fingertips of the right hand within a 7T MRI scanner (see stimulators in *Fig 1A*). The *fingertip localisation task* defined a fingertip region of interest (ROI) for each participant, and the *columnar mapping task* investigated S1 columnar organisation within these ROIs. The two tasks were performed on different days (order counterbalanced) and both involved ∼1 hour of functional data acquisition.

**Fig 1.**
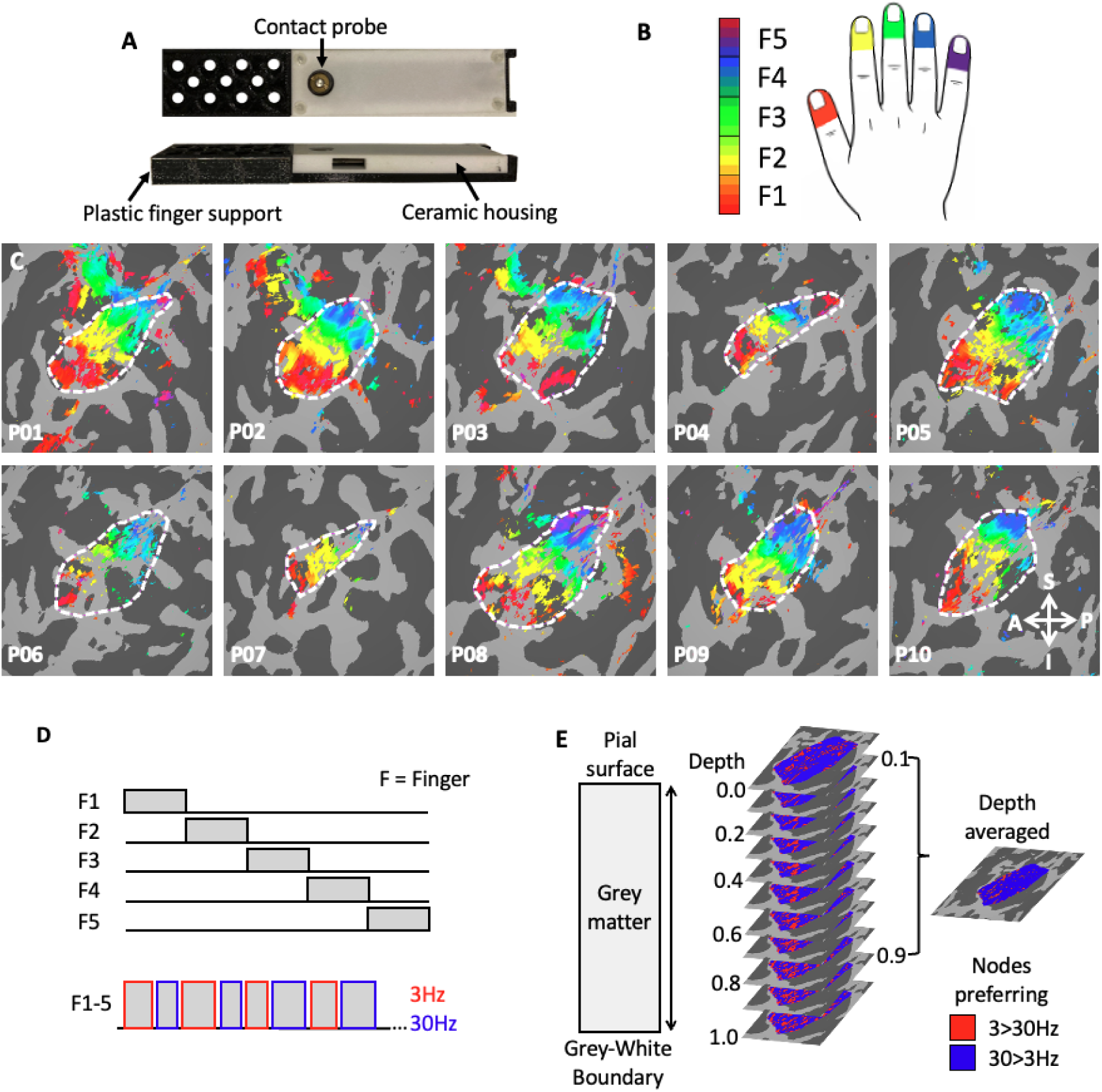
Methods. Two tasks were used to generate preference maps within S1: a columnar mapping task (the experimental task) and a fingertip localisation task (to generate ROIs for the columnar analysis). **A.** Both tasks used stimulation applied to the fingers using magnetic-resonance compatible vibrotactile stimulators (Dancer Designs; Mini-PTS Stimulator System). **B.** In the finger localisation task, the five fingers (F) of the right hand were stimulated in order from 1-5. Colours represent the temporal phase (delay) of the BOLD response relative to the fingertip stimulation cycle. **C.** Fingertip maps for each participant (P01-10). These maps are visualised on the inflated, spherical cortical surface model of each participant’s left hemisphere and are thresholded at q = .001 (FDR corrected; see Fig Supplementary (Supp) 1 for other views and unthresholded maps; see Fig 1B for colour legend). ROIs were manually drawn around the fingertip maps (white dashed line). Anatomical reference coordinates are indicated in the arrow legend at the bottom-right (A = anterior, P = posterior, S = superior, I = inferior). **D.** Stimulation schematic for the two tasks: fingertip localisation task (phase-encoding design; top) and columnar mapping task (alternating 3/30 Hz vibration; bottom). **E.** Depth-averaged analyses averaged activity over cortical depths from the pial surface to the grey-white boundary, excluding the top/bottom 10%, while laminar analyses examined activity at all 11 cortical depths (0.0 to 1.0). Images are from participant P03. For all preference maps, 3 Hz versus 30 Hz preference is indicated by red and blue colours, respectively.

### Fingertip localisation

The fingertip localisation task involved sequential vibration of the five right-hand fingers (F1-F5; *Fig 1B/D*) using a phase-encoded design (Besle et al., 2013; Engel, 2012; Puckett et al., 2017; Sanders et al., 2023). Delay analyses generated fingertip maps (*Fig 1C*), with colours representing the timing of the stimulation sweep. ROIs were manually drawn around the fingertip maps, thresholded at *q* = .001 (FDR corrected; see Methods for ROI criteria).

### Columnar mapping

Within each fingertip ROI, we generated preference maps indicating which cortical regions responded more strongly to 3 Hz versus 30 Hz stimulation. The task involved alternating 6–10s blocks of 3 Hz and 30 Hz vibration applied to all five fingers simultaneously, with counterbalanced start orders and randomised durations (*Fig 1D*). Generalised Linear Model (GLM) regression was conducted on the cortical surface (Dale et al., 1999) and contrasted 3 Hz > 30 Hz activity (Friston et al., 1994).

For the first set of analyses, activity was averaged over the middle 80% (see *Fig 1E*) of the cortical ribbon, removing the top and bottom 10% of cortical volume (Weldon et al., 2019; Wu et al., 2018). This is termed the “depth-averaged” analysis. Depth-averaged preference maps revealed patch-like 3 Hz- (red) and 30 Hz-preferring nodes (blue) for all participants (*Fig 2A; Fig Supp 2*), with every subject showing nodes of both preferences. On average, 76.7% of nodes preferred 30 Hz (range 43.8–96.4%, SEM = 0.04).

**Fig 2.**
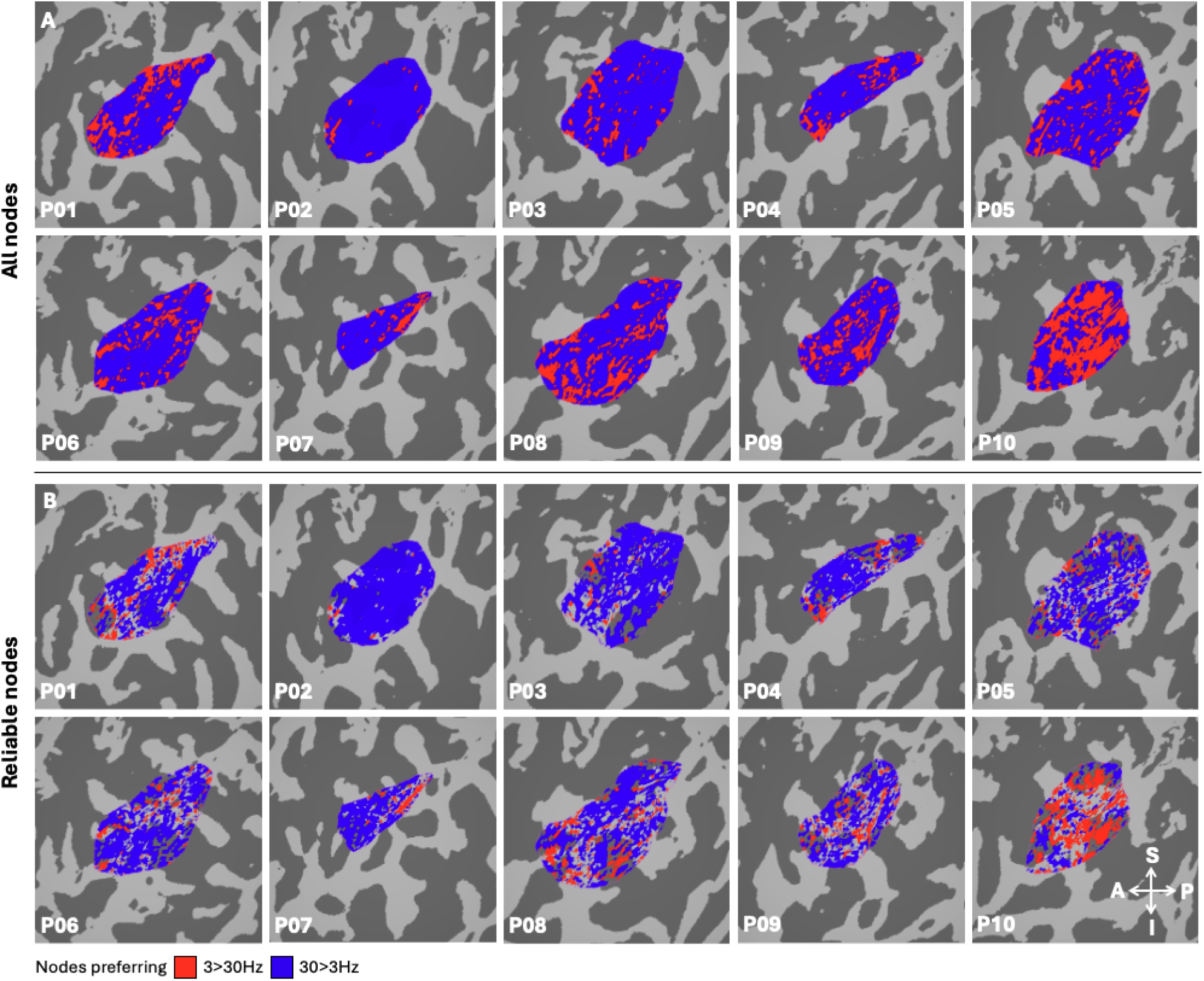
Depth-averaged preference maps for each participant. **A.** Maps generated from the GLM contrast of 3 Hz > 30 Hz. Red/blue colours represent nodes with a preference for 3 Hz/30 Hz stimulation, respectively (colour legend bottom-left). Maps have been binarised for visualisation and are unthresholded. **B.** Maps after removing nodes with unreliable preferences across blocks (split-half analysis), resulting in sparser representations (∼68% of nodes remained, on average). Both panels show depth-averaged data (middle 80% of the cortical ribbon; top/bottom 10% excluded; see Fig 1E), projected onto each participant’s inflated sphere view of the left-hemisphere cortical surface (see Fig Supp 2 for additional views). Anatomical reference coordinates are indicated at the bottom of panel B (A = anterior, P = posterior, S = superior, I = inferior).

### Part 1. Reliability and reproducibility of preference maps in S1

Our first aim was to assess the reliability of the preference maps generated by the 3 Hz > 30 Hz GLM contrast (*Fig 2A*) within our S1 fingertip ROIs. While S1 is expected to reliably encode low-level features of vibrotactile stimulation, some portion of observed reliability could arise from non-specific factors such as statistical noise, global hemodynamic fluctuations, attention, or other imaging-related biases (Polimeni et al., 2018; Wald & Polimeni, 2017). To provide a benchmark for these task-generic effects, we defined a control ROI in frontal cortex. The control ROI was positioned entirely within the partial-volume slab, located in the middle frontal gyrus anterior to the precentral sulcus and superior to the inferior frontal sulcus (see Methods, section *Defining the control ROI*; top of *Fig 3*). While potentially engaged during task performance, this frontal area is not expected to show the same extent of selective preference for 3 Hz versus 30 Hz vibration as S1. As such, it provides a suitable reference for assessing reliability driven by non-specific factors rather than low-level sensory encoding.

**Fig 3.**
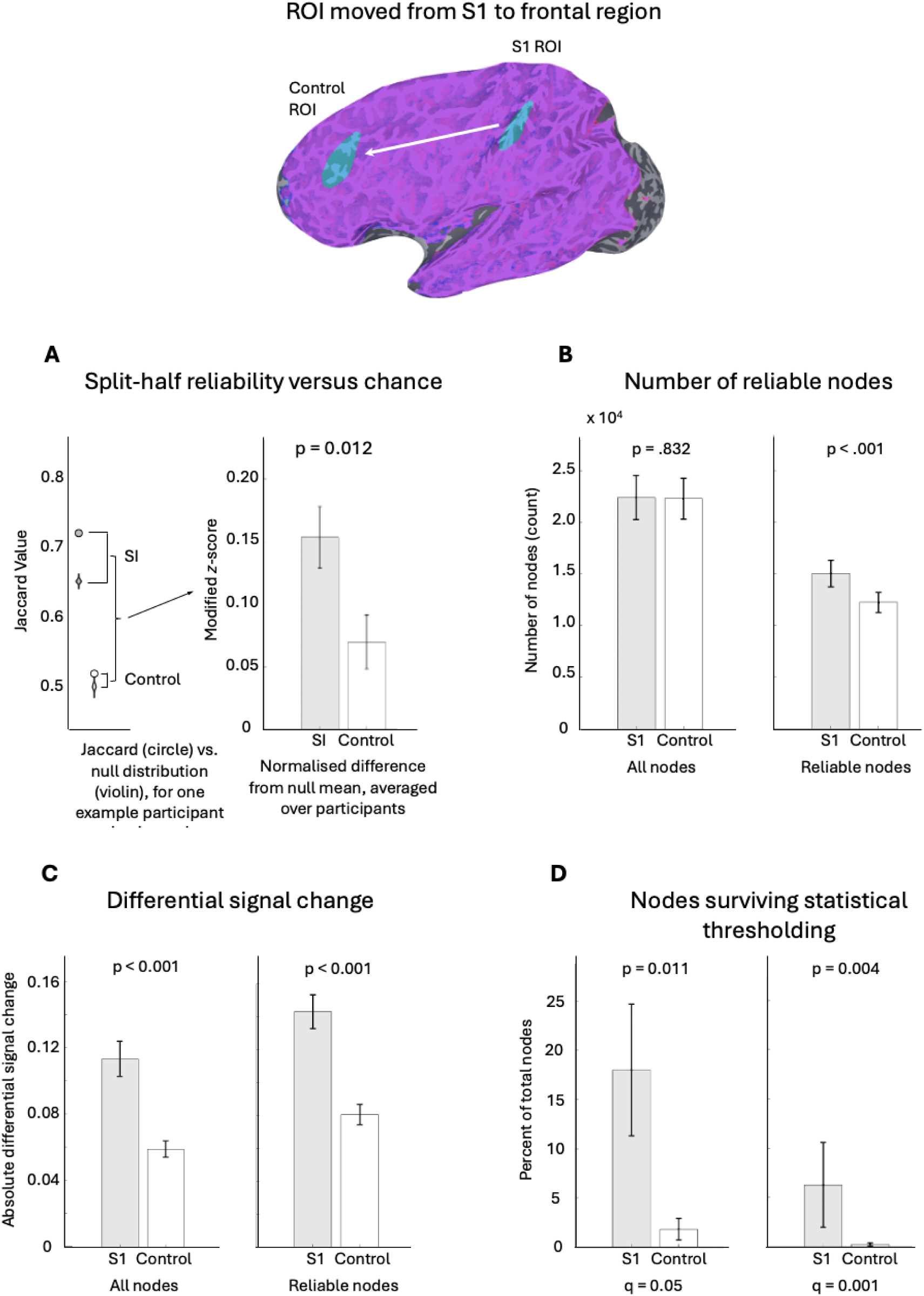
Reliability and reproducibility of preference maps in S1 (grey) and a frontal control ROI of similar size (white). Top panel illustrates how our S1 ROI was shifted on an inflated spherical model of the cortical surface to a manually selected point in the frontal lobe. Purple area shows the imaging slab coverage for a sample participant. Panels **A-D** present results from the four analyses comparing these two regions. **A.** Split-half reliability versus chance: Left panel shows example data for one participant, illustrating the observed Jaccard index (circle) compared with the null distribution from permutation tests (violin). Right panel shows normalised difference from the null mean averaged across participants, representing the difference between observed Jaccard indices and the null distribution. **B.** Number of reliable nodes: Left panel shows the number of all nodes in each ROI (the count). Right panel shows this for reliable nodes only. **C.** Differential signal change (arbitrary units): Left panel shows the distribution of percent signal change difference between 3 Hz and 30 Hz stimulation for all nodes. Right panel shows the same for reliable nodes only. **D.** Nodes surviving statistical thresholding: Left panel shows the count of nodes at q = 0.05. Right panel shows the same at q = 0.001. Error bars indicate SEM.

We anticipated that S1 would show relatively larger differential signal and stronger pattern reproducibility for the 3 Hz > 30 Hz contrast due to its role in encoding vibrotactile frequency. Accordingly, we expected preference maps in S1 would be more reliable across blocks than those in the control region, where reliability would mainly reflect generic contributions (e.g., attention, vascular fluctuations, global signal).

Reliability and reproducibility were compared across ROIs using four analyses: split-half reliability, number of reliable nodes, differential signal change, and nodes surviving thresholding. Paired, Bonferroni-corrected *t*-tests were used for ROI comparisons, except in one case where non-normality of differences violated *t*-test assumptions (*Fig Supp 3*); here, a Wilcoxon Signed-Rank test was applied. For completeness, all paired *t*-tests were repeated with Wilcoxon’s tests, yielding identical results.

#### Split-half reliability versus chance

To assess the reliability of the preference maps, we conducted a within-session split-half analysis, wherein the odd and even blocks of stimulation were compared. The GLM regression was performed separately on the three odd and three even blocks. We then compared the separate odd- and even-block preference maps (i.e., maps from the 3 Hz > 30 Hz contrast). Map similarity was quantified using the Jaccard Index (Jaccard, 1912), which ranges from 0 (no similarity) to 1 (spatially identical; see Methods, section *Split-half reliability analyses* for details). The observed Jaccard Index for S1 and control ROIs is shown in *Fig 3A* for one participant (left panel).

To determine whether this similarity exceeded chance, we generated a null distribution by randomly scrambling 3 Hz and 30 Hz preference at each node separately for the odd and even maps. The Jaccard Index was recalculated for each permutation, repeated 1000 times (Efron & Tibshirani, 1998). The resulting distribution is shown as the violin plot in the left-hand panel of *Fig 3A*.

To compare the observed Jaccard Index to this null distribution, we computed the *normalised difference from the null mean.* This method provides similar information to a *z*-score, while also accounting for observed differences in variance between the S1 and control ROIs (see Methods, section *Split-half reliability analyses* for full rationale and formula). A repeated measures *t*-test confirmed significantly greater normalised difference from the null mean values for S1 compared to the control ROI (*t*(9) = 3.16, *p* = .012; *Fig 3A, right panel*), indicating more consistent 3 Hz/30 Hz preference across blocks in S1. For individual participants, observed Jaccard values lay in the tail of the null distribution in both ROIs, indicating some baseline reliability even in the control region. Note that results were consistent when comparisons were performed using *z*-scores.

#### Number of reliable nodes

We next quantified the number of “reliable” nodes, i.e., nodes showing consistent 3 Hz/30 Hz preference across odd and even blocks. After excluding unreliable nodes, an average of 68.3% of nodes remained (range 54.9% – 89.7%, SEM = 0.03). *Fig 2B* illustrates these masked maps for each participant, showing that reliable nodes were distributed throughout the ROI rather than clustered in specific regions.

Comparing the total number of nodes between S1 and the control ROI revealed no significant difference for all nodes (*t*(9) = 0.22, *p* = .832; *Fig 3B*, left panel), confirming comparable ROI size. In contrast, S1 contained significantly more reliable nodes than the frontal ROI (t(9) = 6.38, *p* < .001; *Fig 3B*, right panel), consistent with its primary role in processing vibrotactile features.

#### Differential signal change

We then assessed the magnitude of differential signal change between 3 Hz and 30 Hz conditions, derived from the 3 Hz >30 Hz GLM contrast. Here, *differential signal change* refers to the difference in estimated BOLD response amplitudes between the 3 Hz and 30 Hz conditions. In S1, the average differential signal increased from 0.11% (SEM = 0.01) across all nodes to 0.14% (SEM = 0.01) when only reliable nodes were considered, indicating that unreliable nodes corresponded to lower signal differences and weaker preference (also see *Fig Supp 4* for further demonstration that unreliable nodes had weaker differential signal).

Comparisons between S1 and the control ROI showed significantly higher differential signal in S1 for all nodes (*t*(9) = 7.99, *p* < .001; Bonferroni-corrected) and for reliable nodes only (*t*(9) = 9.69, *p* < .001; Bonferroni-corrected), supporting the functional specificity of S1 for frequency-selective responses (*Fig 3C*).

#### Nodes surviving statistical thresholding

While our primary analyses focused on split-half reliability to directly assess reproducibility across independent data subsets, we also examined the numbers of nodes surviving statistical thresholding as an alternative reliability measure (*Fig 3D*). Nodes surviving thresholds of *q* = 0.05 and *q* = 0.001 (FDR-corrected) were counted, with the pattern of results supporting the split-half analyses. A repeated-measures *t*-test indicated significantly more nodes survived thresholding at *q* = 0.05 in S1 compared to the frontal control ROI (t(9) = 3.21, *p* = .011), and a Wilcoxon signed-rank test, used for *q* = 0.001 due to violations of *t*-test assumptions (see *Fig Supp 3*), revealed the same pattern (*W* = 45, *p* = .004).

Together, these four analyses converge to demonstrate that preference maps for 3 Hz versus 30 Hz stimulation are more reliable, show stronger differential signal, and survive stricter thresholds in S1 than in the frontal control ROI, consistent with S1’s role in encoding low-level vibrotactile stimuli.

#### Between-session reliability of preference maps

In one participant (33 years old, F), we explored the reproducibility of preference maps generated from scanning sessions separated by 3.5 months (*Fig Supp 5*). Session 2 included 12 blocks of the columnar mapping protocol (total time = 2.75 hours), allowing comparison of within-session reproducibility (odd vs. even blocks) and between-session reproducibility (all 6 blocks of session 1 vs. the first 6 blocks of session 2) with equal power. Between-session Jaccard indices remained well above the null distributions (*Fig Supp 5D*), although the difference between observed and null values appears slightly lower than within-session comparisons (*Fig Supp 5E*).

### Part 2. Laminar analyses

#### Preference pattern across depth

Having established the reliability of depth-averaged preference maps, we next examined whether these preferences were preserved across cortical depth. To examine this, we generated 11 equivolumetric cortical surface models spanning from the pial surface to the grey-white boundary (Polimeni et al., 2010, 2018). Our functional data were interpolated onto these surfaces.

Separate GLMs were estimated at each depth, producing 11 preference maps per participant (see *Fig 1E,* participant P03). Each cortical surface node was classified as preferring 3 Hz or 30 Hz stimulation at each depth, producing a vector of 11 preference values per node (i.e., the *pattern across depth*). This allowed us to assess the consistency of nodes with 3 Hz or 30 Hz preference as identified in the depth-averaged analysis (Part 1). As expected, there was high similarity between the depth-averaged preference maps and the 11 separate depth preference maps (see *Fig Supp 6*).

To quantify depth consistency, for each participant we calculated the percentage of nodes showing each possible depth pattern as a proportion of the total nodes. This was done separately for nodes preferring 3 Hz or 30 Hz in the depth-averaged data. Across participants, the most common pattern for both 3 Hz and 30 Hz preferring nodes was a consistent preference across all 11 depths (*Fig 4A*). On average, only 20% of 3 Hz-preferring nodes exhibited complete depth consistency. In contrast, ∼47% of 30 Hz-preferring nodes showed complete depth consistency (*Fig 4B);* 42.4% when pooling 3 Hz and 30 Hz preferring nodes. The second to fifth most common patterns were similar for both preferences, typically showing a block of consistent preference across most depths (e.g., 7-10 depths) with the opposite preference at either the upper or lower 1-3 depths (*Fig 4A & 4B*). These four patterns accounted for ∼3-6% of the total patterns for either stimulation preference each (∼17% combined).

**Fig 4.**
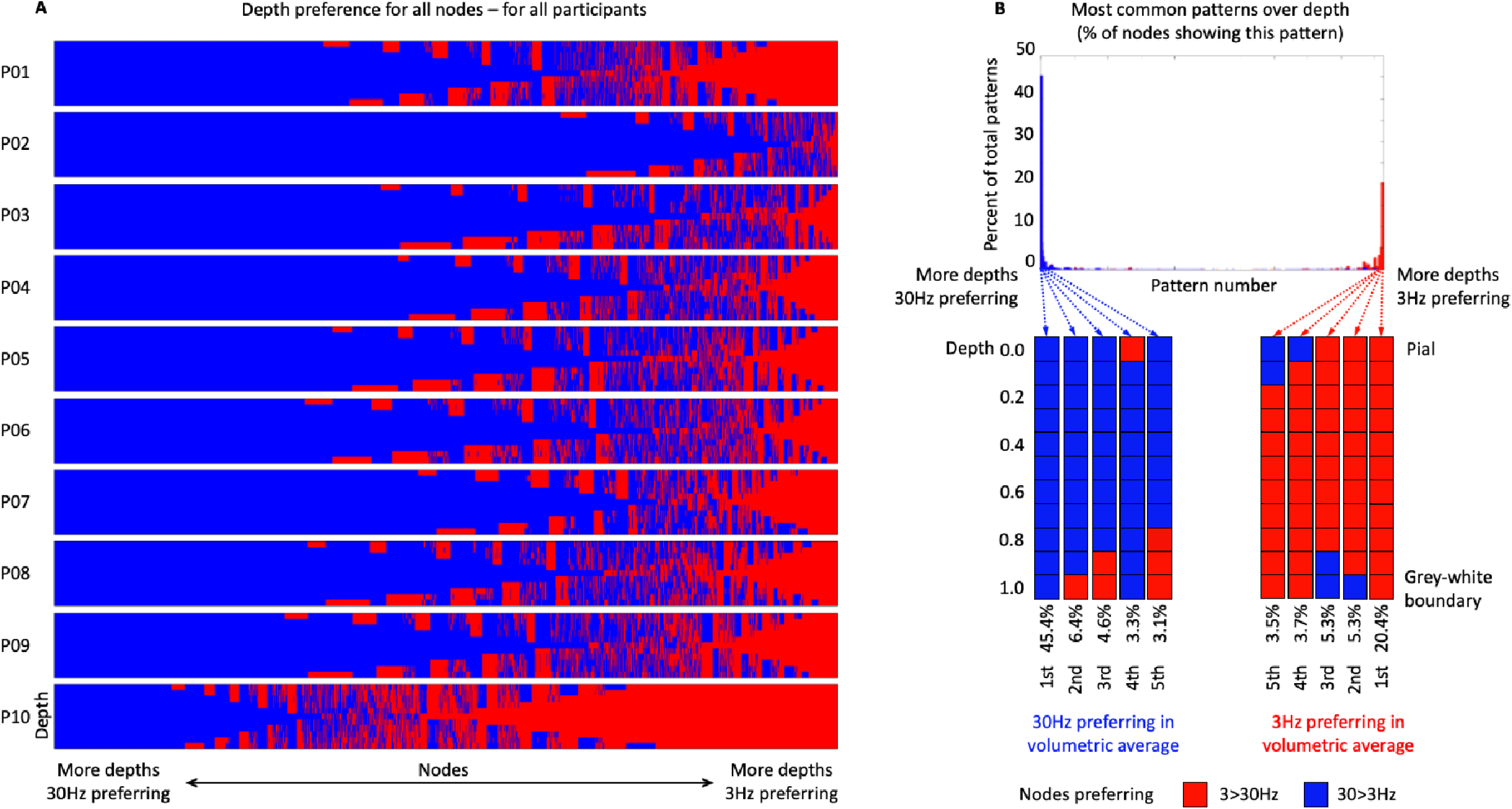
Laminar analyses of depth consistency for vibration preference. **A.** A visual representation of preference for 3 Hz (red) versus 30 Hz (blue) across all 11 cortical depths, for all 10 participants. On each sub-figure, y-axis = depth, x-axis = node number. **B.** Top: Percentage of nodes (y-axis) showing each of the 2,048 possible depth patterns (x-axis), calculated for each participant and then averaged, done separately for nodes preferring 30 Hz or 3 Hz in the depth-averaged data. For example, the far-left (blue) bar shows the proportion of nodes preferring 30 Hz in the depth-averaged data that maintained 30 Hz preference at all depths. Bottom: Red/blue rectangles schematically represent preference at each depth. We show the top 5 most common patterns for nodes preferring 30 Hz (left 5) and preferring 3 Hz (right 5) in the depth-averaged data. Consistent with the concept of cortical columns spanning depth, the most frequent pattern for both 3 Hz and 30 Hz preferring nodes was a consistent preference across all 11 depths, with 30 Hz preferring nodes showing greater depth consistency than 3 Hz preferring nodes.

#### Signal change difference across depth

We next examined the laminar profile of the differential signal (3 Hz > 30 Hz) across all 11 cortical depths for nodes classified by their depth-averaged preference. This analysis characterises how signal strength varies across cortical layers and contextualises occasional deviations from the dominant preference at the most superficial or deepest depths. Percent signal change differences were plotted separately for 3 Hz- and 30 Hz-preferring nodes (*Fig 5A*). When collapsed across depth, 30 Hz-preferring nodes exhibited slightly higher average differential signal (*M* = 0.14%, SEM = 0.01) than 3 Hz-preferring nodes (*M* = 0.10%, SEM = 0.02; paired *t*-test, *t*(9) = 9.83, *p* < .001). This difference appears to decrease when restricting analyses to nodes showing consistent preference across all depths (see *Fig Supp 6B & 6C*).

**Fig 5.**
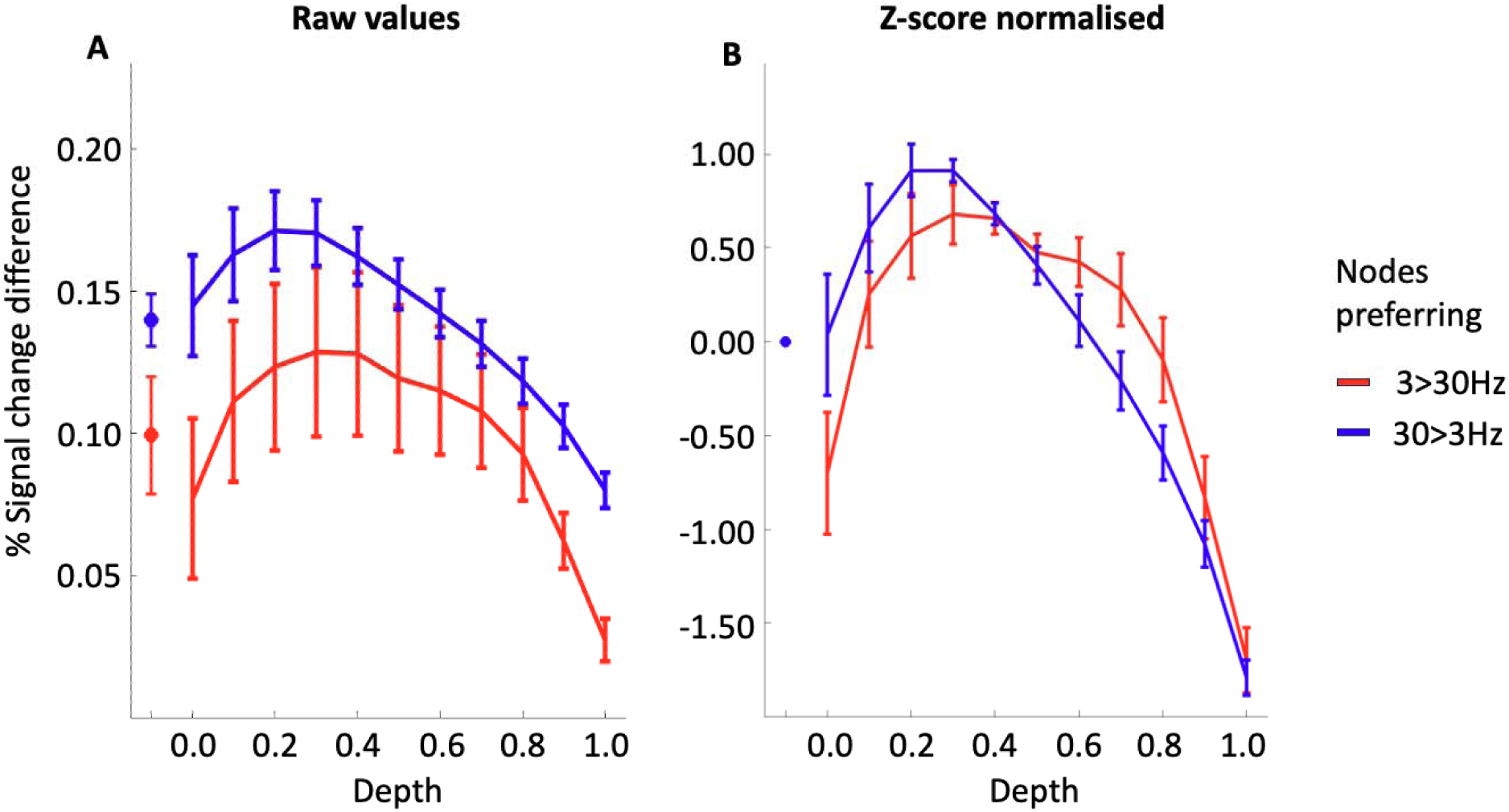
Laminar profiles of differential signal across depth. **A.** Mean percentage signal change difference across depth for 3 Hz (red; mean = 2,584 nodes, SEM = 624) and 30 Hz (blue; mean = 12,455, SEM = 1,491) preferring nodes, classified using depth-averaged data. Data are averaged across all participants; error bars represent SEM. **B.** Same data after z-score transformation within participants to facilitate comparison of profile shapes across depth. In the original data (A), differential signal appears slightly stronger for 30 Hz preferring nodes compared with 3 Hz nodes (dot to the left of each figure represents the mean over depths). The transformed data (B) highlights subtle differences in the shape of the depth profiles (see discussion in main text).

Visual inspection of the raw depth profiles (*Fig 5A*) indicates that both 3 Hz- and 30 Hz-preferring nodes follow a broadly inverse U-shaped pattern, with maximal differential signal at mid-superficial depths and reduced signal at the most superficial and deepest layers. To facilitate comparison of profile shapes across participants, percent-signal-change differences were converted to *z*-scores (*Fig 5B*). Repeated-measures ANOVAs with polynomial contrasts were then applied to quantify whether the depth profiles contained linear (monotonic), quadratic (U-shaped), or cubic (S-shaped) components. For 30 Hz-preferring nodes, all three components were significant (all *p* < .001; .739 ≤ 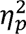 ≤ .756), indicating a combination of overall slope, curvature, and asymmetry across depth. For 3 Hz-preferring nodes, only the quadratic component reached significance (*p* < .001; 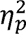 = .92), with non-significant linear (*p* = .084, 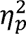 = .30) and cubic components (*p* = .790, 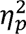 = .01). These results quantitatively confirm the broadly inverse-U depth profiles for both stimulation preferences, while also highlighting subtle differences in their steepness and symmetry.

## Discussion

Our study provides insights into the organisation of the human somatosensory cortex (S1) by identifying patterns consistent with columnar organisation using ultra-high field fMRI. Our first aim was to evaluate the reliability of the preference maps for 3 Hz versus 30 Hz stimulation within S1. Across multiple complementary analyses, we found converging evidence that preference maps in S1 were more reliable and reproducible than those observed in a frontal control ROI. While both regions showed some reliability, likely reflecting task-generic and imaging-related factors, S1 consistently exhibited stronger effects, in accord with its role in representing low-level vibrotactile features. In the one participant tested, preference maps remained reliable even across sessions months apart, highlighting the robustness of these maps despite physiological variability between sessions, e.g., morphometric measures (Nakamura et al., 2015; Trefler et al., 2016) and scanning factors such as head positioning (Raz et al., 2005).

Our laminar analyses revealed that the most common pattern was consistent preference across the entire cortical depth, from the pial surface to the grey-white boundary, a hallmark of columnar organisation (Mountcastle, 1957, 1997; Sur et al., 1981, 1984). The next most prevalent pattern was nodes showing minor preference shifts at superficial or deep layers (∼3–6% of nodes) with these shifts occurring primarily at depths known to be affected by fMRI signal biases, particularly near the grey-white boundary or pial surface (Havlicek & Uludağ, 2020; Polimeni et al., 2010). Taken together, the reliability and depth-consistency results provide evidence consistent with columnar-scale organisation in S1.

While cortical columns are well established in non-human somatosensory cortex (Chen et al., 2001; Dykes et al., 1980; Friedman et al., 2004; Sur et al., 1981, 1984), direct evidence in humans is scarce. Two conference reports have examined frequency-dependent responses across cortical layers, suggesting possible columnar-like patterns (Kim et al., 2021; Yang et al., 2019). However, these studies did not assess reliability or quantify preference strength, which limits interpretability. Our work addresses these gaps by demonstrating reproducible, depth-consistent maps and quantifying differential responses between stimuli.

### Understanding preference in columnar organisation

Approximately 70% of nodes showed a reliable preference for either 3 Hz or 30 Hz stimulation. While this is a majority, many nodes showed weak or inconsistent preference, and even among reliable nodes, differential signals were modest (∼0.14% signal change difference between conditions, collapsed for 3 Hz and 30 Hz preferring nodes). Three factors likely contribute: (1) strong responses to both stimuli, (2) tuning to untested stimulus dimensions, and (3) limits on spatial resolution.

First, it is highly likely many nodes were highly responsive to both stimuli. This is because our ROI comprised voxels showing significant activation to fingertip vibration in an independent localiser task, reflecting genuine somatosensory responsiveness. Within this set, many nodes naturally respond to a range of vibrotactile inputs, consistent with the graded, relative selectivity of S1 columns. Our differential design, with rapid condition switching and minimal inter-stimulus intervals, maximises sensitivity to differences between 3 Hz and 30 Hz stimulation but limits assessment of absolute activation relative to baseline – a trade-off inherent to many columnar mapping studies (Menon et al., 1997; Moon et al., 2007). Future studies employing blocked designs with interleaved rest periods could examine absolute response amplitudes, though at the cost of statistical power for detecting fine-scale columnar organisation.

Second, some nodes may be tuned to stimulus features not tested here. Primate work shows interleaved S1 columns responding to 1Hz, 30 Hz, and 200Hz vibrations (Chen et al., 2001; Friedman et al., 2004), and Mountcastle (1957) described columns preferring skin versus deep stimulation. Low preference may therefore reflect relative rather than absolute selectivity; for example, a “30 Hz-preferring column” may only prefer 30 Hz relative to 3 Hz. Columns may also encode other features such as stimulus amplitude or location, highlighting that “preference” is context-dependent.

Third, technical factors may attenuate observed differences between columns. Our voxel size (∼0.8 mm³) provides high spatial resolution, but it may still be insufficient to target individual cortical columns. Even if a voxel were roughly matched to the size of a column, it is unlikely to be perfectly centred on one, meaning each voxel can sample signals from multiple adjacent columns, which could reduce measured selectivity. Mountcastle (1997) estimated cat S1 columns to be ∼0.3–0.5 mm in width; human S1 columns are likely larger, though precise dimensions remain unknown. Consequently, our voxel resolution may underestimate true columnar selectivity, contributing to the relatively low signal differences observed between nodes preferring 3 Hz vs. 30 Hz stimulation.

### Physiological bases for modest preference

Low selectivity is consistent with neurophysiology. Peripheral mechanoreceptors respond to a range of stimuli, displaying relative rather than absolute selectivity (Saal & Bensmaia, 2014). Slowly adapting mechanoreceptors, for example, respond preferentially to sustained pressure but are also activated by other touch types. In central visual circuits, Hubel and Wiesel (1963) showed that most neurons respond to either eye, exhibiting little or no ocular bias, despite the presence of ocular dominance columns.

High-field human fMRI also reveals sparse columns with low relative preference. Zaretskaya et al. (2020) reported that only 9.6–19% of voxels showed significant ocular preference (at p[<[0.05, uncorrected), resulting in sparse ocular dominance maps. Shmuel et al. (2010) present patches of near-even left/right-eye preference in the visual cortex of humans using 7T fMRI. But, as reviewed by Makin and Krakauer (2023), when examining Schmuel’s Fisher scores, most voxels had very low values, indicating minimal discrimination between eyes. These findings, together with our results, suggest that low relative preference may be a general property of cortical columns and caution against over-interpretation of winner-takes-all maps or simple labelling.

### Conceptualising preference across depth

Our laminar analyses revealed that many nodes maintained consistent preference across all depths (∼20–45%). While fully depth-consistent nodes were the most common pattern, it should be noted they still represented less than half of the total possible patterns. The next four most common patterns (∼3–6% of the total patterns each) showed a preference reversal at one or two depths, typically at the top or bottom.

Importantly, we note that some consistency across cortical depth is expected given the spatial resolution of our data (∼0.8 mm³ isotropic) relative to human S1 thickness (∼2 mm), which spanned roughly 2.5 voxels in our data (Fischl & Dale, 2000). In addition, the presence of draining and pial veins might introduce signal leakage across cortical depths (Polimeni et al., 2010), further reducing the independence of nodes. Indeed, *Fig 5* shows that while there is some increase in 3 Hz > 30 Hz differential signal towards the cortical surface, the maximum signal change is in the upper-middle depths; specifically, at the 5^th^ and 4^th^ depth (of 11) for the 3 Hz and 30 Hz preferring nodes, respectively. Given the maxima are not at the most superficial depths, this suggests that the signal at each node is not overwhelmingly driven by draining and pial vein effects (Polimeni et al., 2010; Uludag & Havlicek, 2021).

The reductions in differential signal at the pial surface and grey–white boundary align with known properties of GRE-BOLD fMRI, where signal loss near the grey–white boundary and surface blurring (Havlicek & Uludağ, 2020; Polimeni et al., 2010) could reduce classification accuracy and produce spurious preference shifts at deep and superficial layers. Since our analyses are performed on the contrast between two tasks, the non-specific effects of veins and fMRI signal biases may be considerably diminished (Havlicek & Uludağ, 2020).

Interestingly, the greater prevalence of fully depth-consistent 30 Hz-preferring nodes (45.4%) relative to 3 Hz preferring nodes (20.4%) is somewhat consistent with findings from previous work. Specifically, as detailed in the introduction, electrophysiology suggests rapidly adapting columns (e.g., 30 Hz preferring) extend over all layers, where slowly adapting columns (e.g., 3 Hz preferring) span only middle layers of the cortex (Sur et al., 1984). Preliminary evidence for this has also been presented in humans in the two conference reports described above (Kim et al., 2021; Yang et al., 2019). In sum, this suggests our depth findings accord with physiological properties of S1 cortical columns, supporting their validity.

### Differences in 3 Hz vs. 30 Hz representation

Across participants, more nodes preferred 30 Hz (∼77%) than 3 Hz (∼23%). 30 Hz-preferring nodes also showed greater depth-consistency (∼45% vs. 20%) and larger differential signals (*Fig 5*). These differences likely reflect the physical and perceptual properties of the stimuli: 30 Hz vibration delivers approximately ten times as many stimulus pulses per unit time and is perceived as more intense (Verrillo et al., 1969). However, a proportion of the 3 Hz-preferring nodes that did emerge showed clear, depth-consistent patterns, indicating that frequency-specific preference can be resolved for both conditions even when stimulus drive differs substantially. We note that the predominance of 30 Hz nodes was only seen in S1, where the split of preferences was similar in our frontal control ROI (see *Fig Supp 7*, bottom), suggesting that the imbalance arises from low-level feature processing rather than other factors. Future work could manipulate vibration intensity independently of frequency to disentangle perceptual and physiological contributions to columnar representation.

### Conclusion

Our study provides evidence for columnar organisation in human S1. We demonstrate reliable, depth-consistent preference maps for two vibration frequencies, with modest but physiologically meaningful selectivity. These findings suggest that S1 columns encode relative preferences rather than absolute stimulus categories, consistent with peripheral and central sensory physiology. Our work establishes a framework for future investigations into the functional properties, perceptual correlates, and stimulus-dependent variability of human cortical columns, with implications for understanding sensory coding across species.

## Ethics approval statement

This project received ethics approval, granted through the University of Queensland Human Research Ethics Committee (reference code: 2021_HE000034).

## Acknowledgements

We thank Thomas Wright for his tireless and scrupulous work in designing and piloting for this experiment.

We sincerely thank Aiman Al Najjar, Nicole Atcheson, and Sarah Daniel for their invaluable assistance in operating the fMRI scanner and for their technical expertise on the project.

The authors acknowledge the facilities and scientific and technical assistance of the National Imaging Facility, a National Collaborative Research Infrastructure Strategy (NCRIS) capability, at the Centre for Advanced Imaging, The University of Queensland.

## Methods

### Participants

Eleven participants were recruited for the experiment. One participant was excluded as only five, rather than six, blocks of the columnar mapping task were collected due to scanner malfunction. This resulted in a final sample of ten participants (7 female, 3 male; *M* age = 26.2 years, SEM = 1.84). Written informed consent was obtained from all participants. Ethical approval was granted by The University of Queensland Human Research Ethics Committee (project identifier: 2021/HE000034).

### General overview

Participants completed two MRI sessions on separate days using a MAGNETOM 7T scanner (Siemens Healthcare, Erlangen, Germany) equipped with a 32-channel head coil (Nova Medical, Wilmington, US). One session involved an independent fingertip localiser task to identify the right-hand fingertip region of interest (ROI) in the left primary somatosensory cortex. The other session consisted of the experimental task used to assess columnar organisation within the localiser-defined ROI.

Each session lasted approximately 1.5 hours. Of this ∼62 minutes were allocated to the acquisition of functional tasks; the remaining time consisted of anatomical imaging, shimming and other preparatory scans. Six participants completed the localiser first, and four completed the columnar mapping task first. All participants completed the two sessions at least 1 week apart (*M* interval = 33.67 days; SEM = 7.98).

### General scanning details

#### Functional Imaging

Functional data were acquired using a 3D gradient-echo echo-planar imaging (3D-EPI) sequence (Poser et al., 2010) with a blipped-CAIPIRINHA implementation (Breuer et al., 2006; Poser et al., 2013; Setsompop et al., 2012). The acquisition slab was tilted to ensure full coverage of left-hemisphere S1, contralateral to the stimulated digits (48 × 192 × 192 voxels, RAI orientation). The sequence parameters were: TR = 1.92 s, TE = 0.0266 s, flip angle = 15°, voxel size = 0.80 × 0.83 × 0.83 mm (RAI), with anterior–posterior phase encoding. GRAPPA was used for in-plane acceleration (reduction factor = 2), and partial Fourier (6/8) was applied. For each run, a total of 320 volumes were collected for the column-mapping task and 195 volumes for the fingertip-mapping task.

At the completion of the main task (columnar or fingertip mapping), a short reverse-phase (i.e., posterior-anterior direction) EPI scan (10 volumes; ∼19 s) was collected for distortion correction (see Methods, section *Pre-processing pipeline – functional data*).

#### Anatomical Imaging

One anatomical image was acquired during both the fingertip localiser and columnar mapping sessions using an MP2RAGE sequence (Marques et al., 2010) with the following parameters: TR = 4300 ms, TE = 2.45 ms, TI1 = 840 ms, TI2 = 2370 ms, FA1 = 5°, FA2 = 6°, voxel size = 0.75 mm^3^. GRAPPA acceleration (in-plane factor = 3) was used. The field of view was ∼192 x 225 x 240 mm with a matrix size of 256 x 300 x 320.

### Vibrotactile stimulation device

Vibrotactile stimulation was delivered to all five fingers of the right-hand using fMRI-compatible piezoelectric stimulators (“Mini-PTS Stimulator System,” Dancer Designs; *Fig 1A*). One stimulator was attached to each fingertip. Each device consisted of a rectangular ceramic housing with a circular opening through which the piezo bender contacted the skin. For stability and ease of attachment, the ceramic housing was embedded in a custom 3D-printed finger sleeve secured using a Velcro strap (also see *Fig 1A*). The stimulators were further secured to the finger using medical tape. The right hand rested on a foam cushion placed at the participant’s midline over the abdomen. Finger spacing was adjusted to prevent contact between stimulators, with positioning maintained by the medical tape.

### Fingertip mapping protocol

#### Stimulation

The fingertip mapping protocol was used to identify the right fingertip ROI in the left hemisphere. A phase-encoded design was employed (Besle et al., 2013; Engel, 2012), with sequential stimulation of the fingers (thumb → little finger = one cycle; see *Fig 1D*). Each finger was stimulated for 9.6 s (5 TRs), with each TR involving a different vibration frequency (5, 20, or 83 Hz; order randomised, no immediate repetitions). Different frequencies were used to reduce adaptation. Responses to specific frequencies were not analysed.

One cycle of stimulation (each of the five fingers) lasted 48 s. Each scanning run contained seven cycles (336 s of stimulation total), plus 19.2 s of baseline at the start and end of the run (10 TRs each). Runs lasted 6.24 min (195 TRs), with each participant completing 10 runs (total scan time = 62.4 min).

#### Task

Participants were instructed to fixate on a central cross during stimulation to help keep their head still. No other task was required.

### Columnar mapping protocol

#### Stimuli

The columnar mapping protocol involved vibrotactile stimulation at 3 and 30 Hz, consistent with prior studies of columnar organisation in non-human primates (Chen et al., 2001; Friedman et al., 2004); for more details see the Introduction.

#### Stimulation protocol

Each scanner run contained two blocks, each with 15 trials per frequency (30 trials/block; 60 trials/run). Frequencies were presented in rapid alternation (3/30/3/30 Hz; see *Fig 1D*), with a 1 TR inter-trial interval. This rapid alternation paradigm was selected to enhance statistical power for contrasting stimulation frequencies in regression analyses (Menon et al., 1997; Moon et al., 2007). Trial durations were 3, 4, or 5 TRs (5.67, 7.68, or 9.6 s), with equal numbers of each duration per block and randomised order. Each run included 7.68 min of stimulation, 1.92 min of inter-trial intervals and 38.4 s of baseline (5 TRs at the start and end of each block), for a total of 10.24 min (320 TRs). Participants completed 6 blocks (total scan time = 61.44 min).

#### Task

To maintain attention, a simple target-detection task was used. During the final 0.33 s of each trial, stimulation ceased for all but one randomly selected “target” finger. Participants indicated the target by pressing the corresponding button on a four-button response box with the fingers of their opposite hand. For trials in which the thumb was the target, participants pressed against the side of the button box. No feedback was provided. In six of the ten participants, accuracy data was missing for one (of six) blocks due to a button-box malfunction. Accuracy was high overall for the four recorded fingers (M = 74.5%, SEM = 3.79).

The target-detection task primarily served to keep participants engaged and ensure they remained awake and alert during stimulation. While attention may have influenced neural responses (Puckett et al., 2017), this was not analysed in the present study. The interaction between attention and columnar organisation remains outside the scope of this work and is left for future investigation.

### Pre-processing general

Unless otherwise specified, pre-processing was performed using the *Analysis of Functional Neuroimages* (*AFNI*) software suite (Cox, 1996) and/or *AFNI’s Surface Mapper* (*SUMA*) program for viewing surface models and mapping volumetric data to the surface (Saad et al., 2004). All steps were conducted separately for each participant.

### Pre-processing pipeline – brain surface models

#### Template generation

An MP2RAGE anatomical image was acquired in both the fingertip localisation and columnar mapping scanner sessions. The two images were combined into a single-subject template using the *antsMultivariateTemplateConstruction2.sh* tool from *ANTS* (*Advanced Normalisation Tools*; https://stnava.github.io/ANTs/; (Shaw et al., 2019; Tustison et al., 2019), creating a balanced template, unbiased toward either scan.

#### Surface reconstruction

The template was submitted to *FreeSurfer’s recon-all* pipeline (https://surfer.nmr.mgh.harvard.edu/fswiki/recon-all) to reconstruct the cortical surfaces. The hires flag was used to preserve native sub-millimetre resolution (Zaretskaya et al., 2018), along with an expert configuration file to ensure proper inflation of the high-resolution cortical surfaces.

#### Skull stripping template

Although *FreeSurfer* automatically produces a skull-stripped image, skull-stripped images often retain dura or skull remnants (Bazin et al., 2014). To improve this, *FreeSurfer* outputs from recon-all autorecon1 were processed with *antsBrainExtraction.sh* (*ANTS)* and then refined using *Segmentator* (https://github.com/ofgulban/segmentator) where possible. The resulting (further) skull-stripped images were resubmitted to *FreeSurfer’s autorecon2* and *autorecon3* steps to regenerate cortical surfaces.

#### Manual cleaning for more accurate surface models

Manual brain mask and segmentation cleaning was performed in *FreeSurfer’s Freeview* for all participants, to resolve issues that occur during the reconstruction process. Cleaning focused on the motor and primary somatosensory cortices, defined by anatomical landmarks (e.g., between the pre- and post-central sulci) and the Destrieux segmentation atlas (Destrieux et al., 2010). This ensured that the independently defined ROIs were fully contained within the cleaned region. Cortical surface models were regenerated following cleaning of the segmentation and manually inspected.

#### Resampling of cortical mesh

The initial cortical mesh matched the anatomical image resolution (0.75 mm isotropic). To prevent data loss during interpolation of functional data (Wang et al., 2022), the mesh was refined using *FreeSurfer’s mris_mesh_*subdivide butterfly subdivision method. Two successive iterations were applied, yielding a final surface mesh with an average inter-node distance of 0.20 mm, SD = 0.09 (original surface reconstruction *M* = 0.79 mm, SD = .38).

#### Laminar surface generation

Using the anatomical template, 11 laminar surfaces were generated per participant (nine intermediate and two boundary surfaces). This number was chosen in line with previous surface-based approaches (Polimeni et al., 2010, 2018). On a practical level, this choice allows for a sampling gradient across depth, rather than the arguably arbitrary bins of superior, middle, and deep, for example. The laminar surfaces were equivolumetric (Waehnert et al., 2014, 2016), and spanned from the pial surface to the grey-white matter boundary (see *Fig 1E*). They were generated with the ‘*Surface Tools*’ package (Wagstyl et al., 2018); https://github.com/kwagstyl/surface_tools). Each surface comprised the same number of nodes to ensure node-wise correspondence across all surfaces.

### Pre-processing pipeline – functional data

Functional data for fingertip localisation and columnar mapping were pre-processed separately, following the steps outlined below.

#### Outlier count

Voxel-wise intensity outliers were identified using *AFNI’s 3dToutcount* (settings: automask, detrend with 5^th^ order Legendre polynomial). This procedure detects voxels whose intensity deviates substantially from neighbouring voxels over time, often reflecting participant movement or scanner artifacts. The fraction of outlier voxels at each time point was saved as a one-dimensional text file for use in the regression analysis (columnar mapping data only; fingertip localisation used a delay analysis instead). Volumes in which >5% of voxels were outliers were censored, as per AFNI default values.

#### De-spiking

Spikes in signal intensity from the voxel time-series were removed using *AFNI’s 3dDespike* (settings: ‘new’ method for computing fit, no mask). Local median filtering identified deviations, which were then replaced via interpolation with neighbouring time-points, preserving the underlying signal structure.

#### Blip up/down distortion correction

Susceptibility-induced distortions in the EPI data were corrected using *AFNI* tools and reverse phase-encoded functional images. Median datasets were computed for forward and reverse time series (*3dTstat*) and masked to isolate brain regions (*3dAutomask*). *AFNI’s 3dQwarp* then calculated midpoint warp fields to align the forward and reverse data, which were applied with *3dNwarpApply* to correct distortions. These warp fields were subsequently applied to the full EPI time series to reduce spatial misalignment caused by magnetic field inhomogeneities, ensuring better correspondence between functional data and underlying anatomy while maintaining image obliquity.

#### Volume registration

Motion correction was performed in two stages: (1) registering volumes within each run, and (2) registering blocks together, using *AFNI’s 3dvolreg* with rigid-body (six-parameter) transformations and Fourier interpolation. In both cases, the base volume was the time point with the lowest intensity outlier fraction. Transformation matrices from all correction steps were combined prior to application to reduce interpolation-related smoothing (Polimeni et al., 2018).

#### Scaling

EPI intensities were scaled to a mean of 100 (range 0-200) using *AFNI’s 3dTstat* and *3dcalc* to avoid negative values and standardize signal across voxels.

#### Alignment

An initial rough alignment of the anatomical template to the EPI base volume (the reference volume used for EPI motion correction) was performed manually using ITK-SNAP (Yushkevich et al., 2006). Because the EPI reference typically covers only the imaged slab rather than the whole brain, this step produced a slab-limited anatomical image and an approximate transform. This preliminary alignment provides a stable starting point and helps prevent failures during more precise whole-brain registration.

Final alignment was achieved with *AFNI’s 3dAllineate* (cost function: lpc+), which used the preliminary transform as the initial estimate for aligning the full anatomical volume to the EPI base. Automasking, autoweighting, a two-pass strategy, and wsinc5 interpolation were applied. This process yielded a fully aligned whole-brain anatomical image optimized for accurate registration with the functional EPI data. Manual inspection was conducted to verify effective alignment.

#### Generation of SUMA surface files

Specification files for *SUMA* visualization were generated using a modified version of *AFNI’s SUMA_make_spec_FS*, allowing inclusion of intermediate surface models, output in GIFTI format, and inflation over 15,000 to accommodate the high-density cortical mesh.

#### Alignment of surface model to anatomical template

The alignment of the surface model to the anatomical template was then performed to ensure accurate registration for subsequent analyses. The surface model anatomical image was first roughly aligned to the template using *AFNI’s 3dAllineate* (cost function: lpa, wsinc5, automask/autoweight, two-pass, center-of-mass alignment). Final alignment was performed with *SUMA_AlignToExperiment*, using the previous rough alignment as the starting transform. The -wd flag allowed use of 12-parameter affine transformations via *3dWarpDrive.* The final registration was visually inspected before post-processing was conducted.

### Post-processing pipeline – fingertip maps

#### Trimming and averaging

The baseline TRs at the beginning and end of the EPI time-series were removed to retain only the stimulation (phasic) portion of the data, i.e., rest / baseline was removed. An average time-series was then calculated across all ten blocks of the fingertip mapping EPI data.

#### Resampling of functional data

Prior to surface-based sampling, the averaged functional volumes were upsampled to a finer grid spacing (0.4 mm isotropic from ∼0.8 mm isotropic) using AFNI’s *3dresample* with nearest-neighbour interpolation (default settings). This upsampling reduces interpolation-related spatial blur when sampling functional data at arbitrary coordinates along cortical depth trajectories, as the spatial extent of interpolation kernels scales with voxel grid spacing (Wang et al., 2022; Polimeni et al., 2018). Additionally, finer grid spacing provides improved discretisation of the continuous depth profile for laminar analysis and reduces the number of cortical voxels ‘missed’ during surface projection due to mesh-voxel grid misalignment.

#### Interpolation of functional data to surface model

Next, we mapped the volumetric EPI data onto the surface model of the brain for each participant (see Methods, section *Pre-processing pipeline – brain surface models*). In sum, we took the middle 80% of our functional data in between our pial and grey-white boundary surface models (top and bottom 10% removed) and averaged the data in-between. This created one value that was interpolated onto the pial surface model. This dataset is referred to as the ‘depth-averaged’ data to differentiate it from the data used for our laminar analyses.

Excluding the top and bottom depths, as done here for the depth-averaged data, is common practice in laminar imaging to reduce the effect of surface blurring at superficial depths and signal-to-noise drop-out at deep depths (Scheeringa et al., 2023). However, in the laminar analyses we retained these depths to examine how such biases might affect the results (see *Discussion*).

To interpolate our functional data to the surface, for our depth averaged analyses, we used *AFNI’s 3dvol2surf* (settings: map from layer 10 (A) to layer 0 (B); an ‘average’ mapping function (filter for values along the segment), i.e., output the average of all voxel values along the segment; use all segment point values in the filter (‘nodes’ index); 10 evenly spaced steps along each segment; the grid parent was the resampled averaged EPI data from the previous step). For the laminar analyses, the ‘mask’ (not average) setting was used to only extract values intersecting the nodes of the surface.

#### Delay analysis

To assess the temporal dynamics of the neural response to fingertip stimulation, we performed a delay analysis on the averaged volumetric EPI data from the fingertip mapping blocks. This analysis was conducted using *AFNI’s 3ddelay*, which allows for the identification of temporal shifts between the voxel time series and a reference waveform. We used a sinusoidal reference wave (Saad et al., 2003) and the default settings for *3ddelay*.

For each voxel, the delay between its time series and the reference time series was calculated, with each voxel then assigned a colour representing this delay. The delay values were interpreted with respect to the timing of fingertip stimulation. These fingertip maps are illustrated in *Fig 1C* and the colour legend in *Fig 1B*. A delay value of five, for instance, indicates that the voxel’s time series is delayed by five seconds relative to the reference time series.

#### Generation of fingertip ROIs

Fingertip maps were thresholded using the correlation between the time series and the reference waveform, controlling the false discovery rate at *q* = .001 (Genovese et al., 2002). This threshold is more conservative than the commonly used *q* = .05 and was chosen to minimise inclusion of sensory-driven responses in adjacent motor cortex (Genovese et al., 2002; Puckett et al., 2017).

The ROIs were manually drawn on the fingertip maps by an experienced member of the research team with expertise in somatotopic mapping. A set of guidelines was developed to guide this process, in conjunction with the entire research team. The aim was to ensure that the significant activation was closely and smoothly enclosed, avoiding sharp deviations in the trajectory of ROI boundary line. These ROIs were visually inspected and agreed upon by two other team members with somatotopic mapping expertise. Visual inspection, in relation to major anatomical landmarks, indicated that the ROIs are primarily located towards the anterior bank of the post-central gyrus (see *Fig Supp 1*). This may approximately correspond with Brodmann Area 3b, though the exact location is difficult to definitively determine without histology.

**Post-processing pipeline – columnar maps** *Resampling & interpolation of functional data* As in fingertip mapping protocol above.

#### Regression

To probe for columnar organisation, we conducted an ordinary least-squares regression on the column time-series data within the fingertip region ROI. Specifically, we use *AFNI’s 3dDeconvolve*; settings: mask using ROI, censor volumes using motion file (generated in *Outcount* block), detrend the baseline using 1^st^-5^th^ degree polynomials, dmblock for duration modulated blocks (as our trials for 3/30 Hz ranged from ∼6-10 seconds see Methods, section *Columns protocol*). We also specified one contrast in the model: 3 Hz > 30 Hz.

### Post-processing pipeline – laminar columnar maps

For each run of our EPI data, the volumetric data was interpolated separately onto the 11 surface models spanning from the pial surface to the grey-white boundary using the methods described above, using the -map_func mask option in AFNI’s *3dVol2Surf*. This was to ensure that only the EPI data intersecting the respective surface model was mapped). Please note, the volumetric data described here is the same functional EPI data described above for the columns depth-averaged maps, therefore, the pre-processing pipeline is also detailed above.

We then conducted a separate regression for each depth (using the same method described for the depth-averaged data, i.e., using *3dDeconvolve*, with analysis confined to the ROI). This approach allowed us to probe the organisation of columnar patterns at different depths within the cortex, by yielding a set of preference maps (3/30 Hz) for each node.

### Defining the control ROI

In Part 1 of our analyses, we compared reliability and reproducibility between our S1 ROI and a frontal, control ROI. To do so, we moved our S1 ROI to the frontal cortex. This allowed us to largely preserve the shape, size and number of nodes of the ROIs between regions.

To perform the move, we extracted the Cartesian (RAI) coordinates for the nodes in the S1 ROI and computed the centroid of this region. Using AFNI’s *Surf2VolCoord,* we identified the node within the S1 ROI closest to this Cartesian centre point. We then manually selected a target point in the frontal lobe to which the ROI would be translated. This location fell within the lateral prefrontal cortex and likely corresponds approximately to dorsolateral prefrontal cortex in most participants (see top of *Fig 3* for example in a representative participant). The translation vector between the S1 centroid and this frontal target point was computed in RAI coordinates and applied that translation to the coordinates of all nodes in our S1 ROI. The node numbers on the cortical surface model closest to the translated coordinates were then identified. Duplicate node numbers (arising when multiple translated coordinates mapped to the same node) were removed. Because this slightly reduced the size of the moved ROI, the resulting frontal ROI was dilated by one iteration using AFNI’s 3dmask_tool.

### Higher-order analyses

#### Split-half reliability analyses

To assess the reliability of our preference maps, we performed a within-session, split-half reliability analysis. Specifically, the six total blocks were divided into two subsets: the three odd-numbered blocks and the three even-numbered blocks. The GLM analysis used to generate preference maps (based on the 3 Hz > 30 Hz contrast) was applied separately to each subset. The resulting preference maps from the odd and even blocks were then compared to evaluate their similarity.

The Jaccard Index (Jaccard, 1912) was used to quantify the spatial similarity between the two maps. The Jaccard Index is a measure of overlap, ranging from 0 (no overlap) to 1 (perfect overlap). It is calculated as:

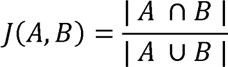

where *A* and *B* represent the sets of active regions in the two maps, ∣ *A* ∩ *B* ∣ is the size of the intersection of the sets, and ∣ A ∪ *B* ∣ is the size of their union.

The same procedure was used for our between-sessions analyses, but with different subsets of the data collected (see *Fig Supp 5*).

To determine whether the observed similarity between odd- and even-run preference maps was greater than expected by chance, we generated null distributions of Jaccard Indices for each participant. To create these distributions, the 3 Hz > 30 Hz preference values at each node were randomly permuted separately for the odd and even maps. Jaccard Indices were then recalculated for the scrambled maps. This process was repeated 1000 times using a bootstrapping procedure to generate a null distribution of chance-level Jaccard Indices for each participant (see example from a representative participant in *Fig 1A*).

For each participant, we quantified how far the observed Jaccard Index deviated from their participant-specific null distribution using a normalised difference metric, here called *normalised difference from the null mean*. Specifically, we scaled the difference between the observed Jaccard index and the null mean by the distance from the null mean to the theoretical upper bound of the Jaccard index (1):

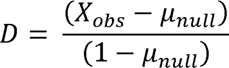

where X_obs_ is the observed Jaccard index and µ_null_is the mean of the null distribution. Larger values of *D* indicate greater similarity between odd and even blocks than expected under the null.

This approach was chosen instead of conventional *z*-scoring because *z*-scores normalise the observed deviation by the standard deviation of the null distribution. In our data, the S1 ROI contained a substantially higher proportion of 30 Hz-preferring nodes (∼77%) than 3 Hz-preferring nodes, introducing a structural class imbalance that inflated the variance of the null distribution independently of true odd–even similarity. This 3 Hz versus 30 Hz imbalance was not seen in the control ROI (see *Fig Supp 7*). As a result, *z*-scores would disproportionately penalise the condition with greater representational heterogeneity, driven by node prevalence rather than signal reliability. Normalising by the distance to the theoretical maximum avoids variance-based scaling and provides a bounded, interpretable measure of effect magnitude relative to chance.

The same procedure for computing Jaccard indices and generating participant-specific null distributions was applied to between-session analyses, using data subsets drawn from different sessions.

#### Laminar analyses – Preference pattern across depth

For each participant, we quantified the number of cortical nodes exhibiting each possible preference pattern across cortical depths. For example, we calculated how many nodes consistently preferred 30 Hz at all depths. There were 2048 possible combinations of 3 Hz and 30 Hz preferences across 11 depths.

For each participant, the proportion of nodes exhibiting each of the 2048 patterns was calculated. To do so, we divided the number of nodes showing a given pattern by the total number of nodes for that participant. This yielded the percentage of total nodes displaying each pattern. These percentages were then averaged across participants. This analysis was conducted separately for nodes that preferred 30 Hz in the depth-averaged data and those that preferred 3 Hz in the depth-averaged data.

#### z-score transformation of signal change difference across depth

To aid comparison of the distributions of signal change difference across depth, the signal change difference values for each participant were transformed into *z*-scores, and the absolute values were taken (*Fig 5B*). This was done *to* standardise the distributions by centring them at a mean of 0, thereby controlling for differences in the overall magnitude of signal change difference. Additionally, *z*-scoring adjusted for differences in variance between distributions (such as may be exist for nodes preferring 3 Hz versus 30 Hz; see error bars in *Fig 5A*) by scaling the standard deviation of both distributions to 1.

The absolute value of the *z*-score, ∣*z*∣, was used to quantify the magnitude of the standardised deviation while disregarding direction (resulting as a function of our 3 Hz > 30 Hz contrast, where 30 Hz preferring nodes have negative signal change difference values).

