## Supplementary Figures for "Ultra-high field fMRI reveals functional patterns consistent with columnar organisation in human somatosensory cortex"

**Supplementary Materials**

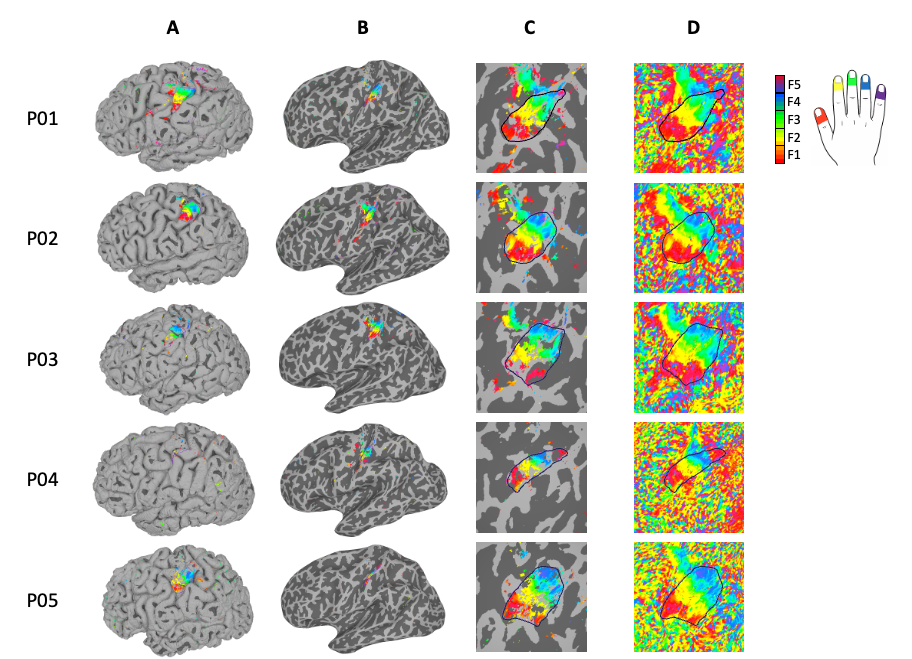

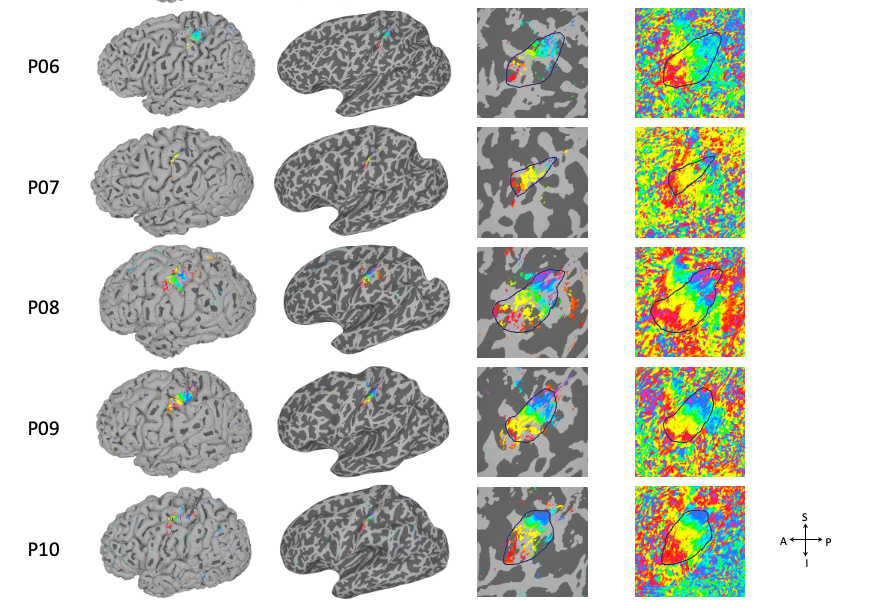

***Fig Supp 1.*** *Fingertip maps for each participant (P01-10), with additional views to those shown in the main text. The maps contain a region of interest (ROI) that was manually drawn around the fingertip representations (dark purple line). All maps are visualised on the left-hemisphere cortical surface. Columns A-C have been thresholded at q = .001 (FDR corrected). Column D shows the unthresholded maps. The maps differ in the inflation of the cortical surface:* ***A****. Uninflated ‘pial’ view,* ***B****. Semi-inflated view,* ***C & D.*** *Fully inflated ‘spherical’ view (as presented in the main text figures). See the top right for the colour legend. For anatomical reference coordinates see the arrow legend at the bottom-right of panel D (A = anterior, P = posterior, S = superior, I = inferior).*

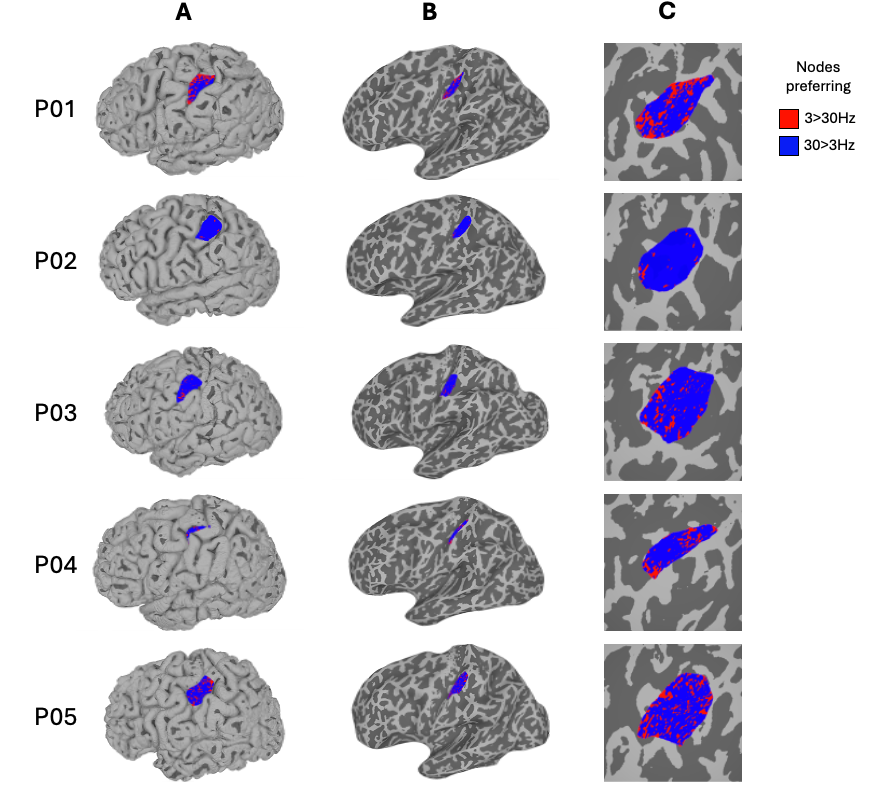

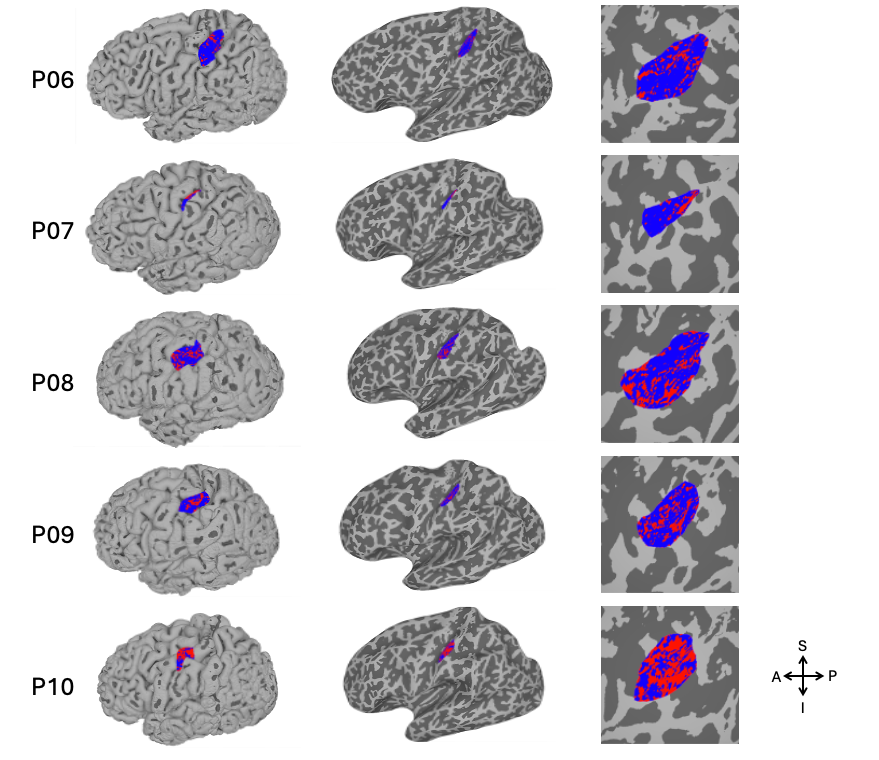

***Fig Supp 2.*** *Preference maps for each participant (P01-P10) generated by the GLM contrast of 3 Hz > 30 Hz. Maps differ in the inflation of the cortical surface:* ***A.*** *Uninflated ‘pial’ view.* ***B.*** *Semi-inflated view.* ***C.*** *Fully inflated ‘spherical’ view (as presented in the main text figures, e.g., Fig 2A & 2B). The maps presented are unthresholded. Red/blue colours represent nodes with a preference for 3 Hz/30 Hz stimulation, respectively (see the colour legend at top-right). All maps use depth-averaged data (activity averaged over the cortical depth, with the top/bottom 10% removed). The maps were binarized for visualisation purposes. The left hemisphere of the cortical surface is presented (right hand stimulation). For anatomical reference coordinates see the arrow legend at the bottom right. Please note: A = anterior, P = posterior, S = superior, I = inferior.*

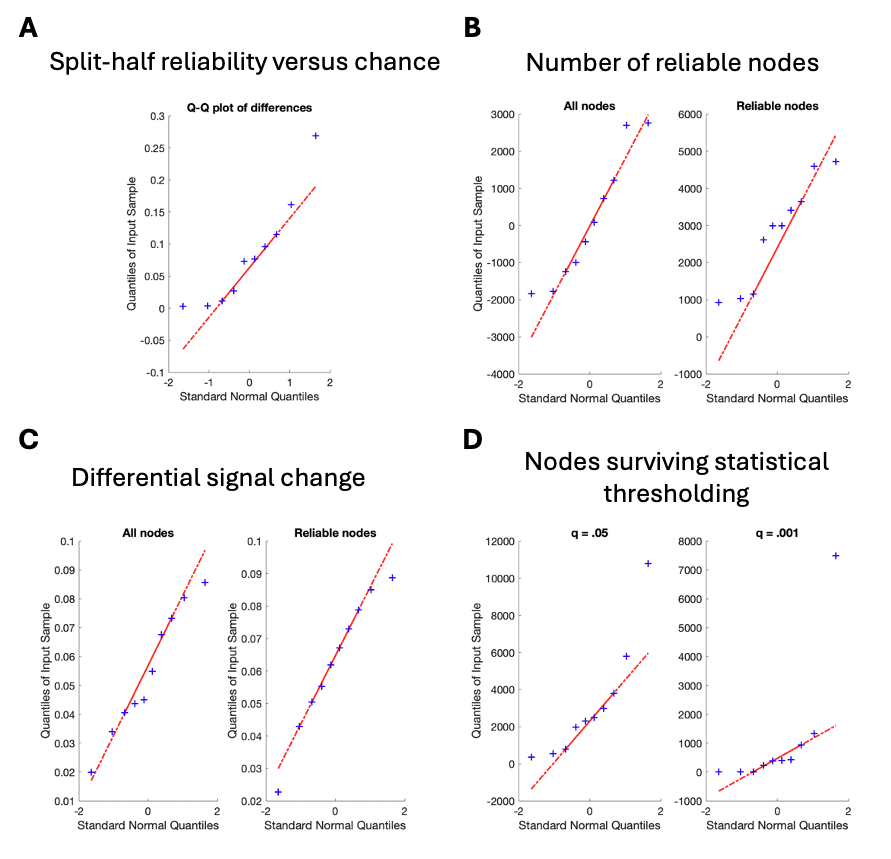

***Fig Supp 3.*** *Q-Q plots used, in conjunction with Kolmogorov-Smirnov tests presented in main text, to assess violations of assumptions of t-tests. Specifically, to investigate non-normality of the distribution of the differences being tested. The four panels (A-D) correspond with the four analyses presented in Part 1 of the main text.*

**Tests for assumption violations for reliability and reproducibility (Part 1) t-tests**

We conducted Kolmogorov–Smirnov tests and inspected Q-Q plots to test for violations of the assumptions of parametric *t*-tests. Specifically, we used these tests to probe for non-normal distribution of the differences between values being tested. For the comparisons in *Fig 3A*: Split-half reliability versus chance, *Fig 3B*: Number of reliable nodes, and *Fig 3C*: Differential signal change, the Kolmogorov–Smirnov tests were not significant (all p ≥ .832) and the Q-Q plots suggested no issues. This suggests the assumptions of parametric *t*-tests were not violated, and thus, supported their use for these analyses.

For our comparisons in *Fig 3D*: nodes surviving statistical thresholding, the tests for assumption violations were also non-significant for comparisons at *q* = .05: Kolmogorov-Smirnov (*p* = .612), and *q* = .001 (*p* = .107). For the *q* = .05 comparisons, however, the Q-Q plot revealed an outlier in the distribution of differences (see *Fig Supp 3D*, right panel above). In the interests of caution, we used a non-parametric, Wilcoxon’s signed rank test for this one comparison. For completeness, we also repeated all our parametric t-tests with Wilcoxon’s. The results were identical.

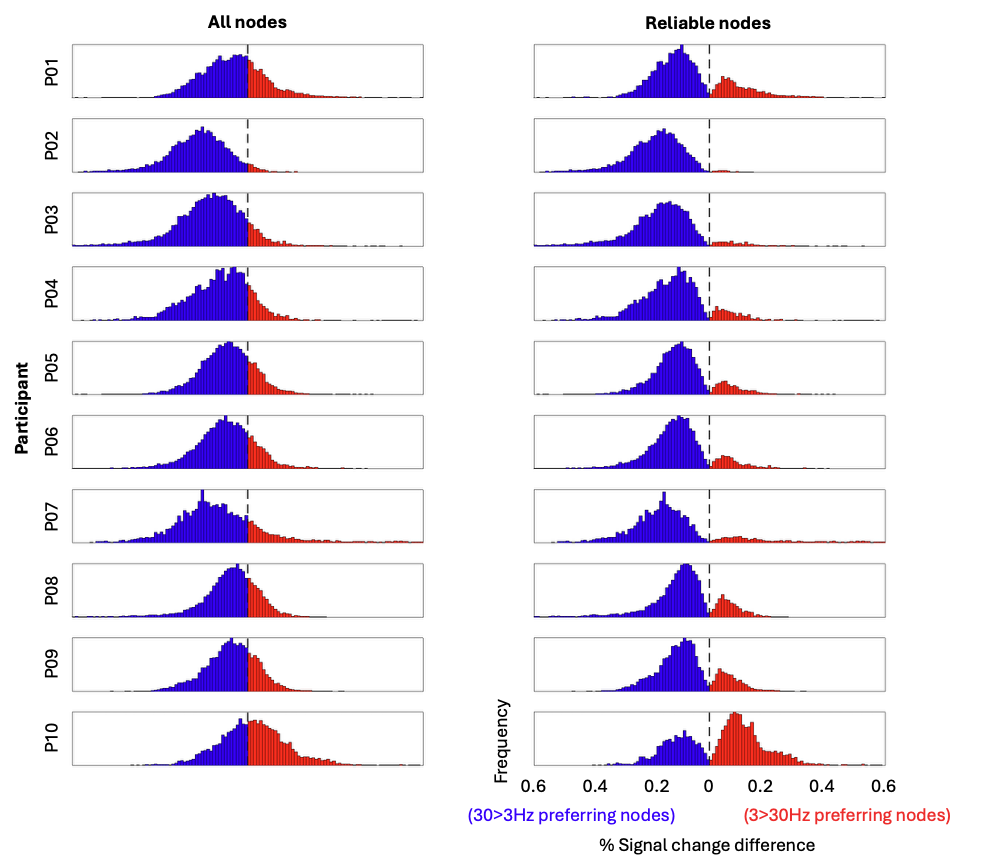
***Fig Supp 4.*** *Histogram of 3 Hz > 30 Hz differential signal change values for each of the ten participants (different rows). On the x-axes, zero represents no differential signal difference between 3 Hz versus 30 Hz vibration stimuli. Blue bars represent signal difference for 30 Hz > 3Hz and red bars represent 3 Hz > 30 Hz. The x-axis values were truncated (the long tails beyond ~6 standard deviations of some distributions removed) for visualisation purposes. Y-axes show the frequency of differential signal change values. The y-axis values are not presented for each figure as they depend on the total number of nodes in the region of interest and are therefore largely arbitrary (though all axes start at 0 and range up to either 500 or 1000). The left panel shows differential signal change values for all nodes, whereas the right panel shows reliable nodes only.*

As can be seen in *Fig Supp 4*, removing unreliable nodes had the effect of removing many nodes with a low differential signal change difference between the 3 Hz and 30 Hz conditions (i.e., from the GLM 3 Hz > 30 Hz contrast). The unreliable nodes were predominantly the near zero values. Thus, removing them led to a more bimodal distribution (see *Fig Supp 4*, right panel).

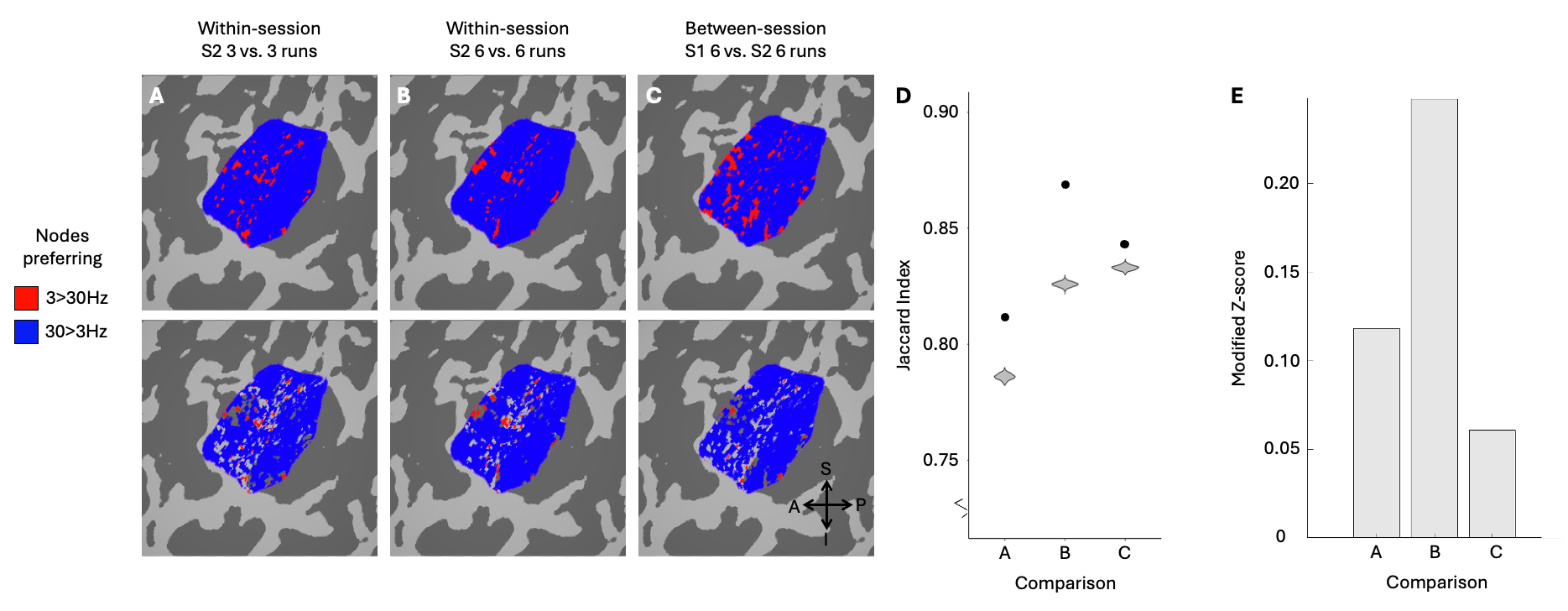

***Fig Supp 5.*** *Between-session reliability of preference maps for one participant who underwent two sessions (P03). Please note: This participant completed 12 runs in the second session (instead of six, as for all other sessions/participants) to allow for additional analyses. Red/blue colours represent nodes with a preference for 3 Hz > 30 Hz/30 Hz > 3 Hz stimulation, respectively (see the colour legend on the right). Note: maps have been binarized for visualisation purposes.* ***A.*** *Top: Preference map for P03 session two, runs 1-6 (i.e., 6 runs, as for preference maps in main text Fig 2). Bottom: Preference map from A (top) with nodes that were inconsistent between the 3 odd runs and 3 even runs (runs 1-6).* ***B.*** *Top: Preference map for P03 session two, runs 1-12. Bottom: Preference map from B (top) with nodes removed that were inconsistent between the 6 odd and 6 even runs (runs 1-12).* ***C.*** *Top: Preference map for P03 session one, run 1-6 (also seen in Fig 2A). Bottom: Preference map from C (top) with nodes removed that were inconsistent between the 6 runs of session one and 6 runs of session two. All maps use depth-averaged data (activity averaged over the cortical depth, with the top/bottom 10% removed). The maps are visualised on an inflated sphere view of the left hemisphere cortical surface, and are statistically unthresholded. For anatomical reference co-ordinates see legend at the bottom of C (A = anterior, P = posterior, S = superior, I = inferior).* ***D****. Observed Jaccard indices (circles) and the null distribution of Jaccard indices generated from scrambled maps (violins).* ***E.*** *Normalised difference from the null mean values, representing the distance of the observed Jaccard value from the null distribution (see Methods, section Split-half reliability analyses).*

As discussed in the main text, the Jaccard index calculated on between-session data (*Fig Supp 5C*) sat well outside the null distribution, suggesting it was more reliable than would be expected by chance. This is similar to what was found in the within-session data (e.g., *Fig Supp 5A & 5B*). As expected, within-session the distance was larger when more data were included (6 runs per session; Fig Supp 5B) compared with fewer runs (3 runs; Fig Supp 5A).

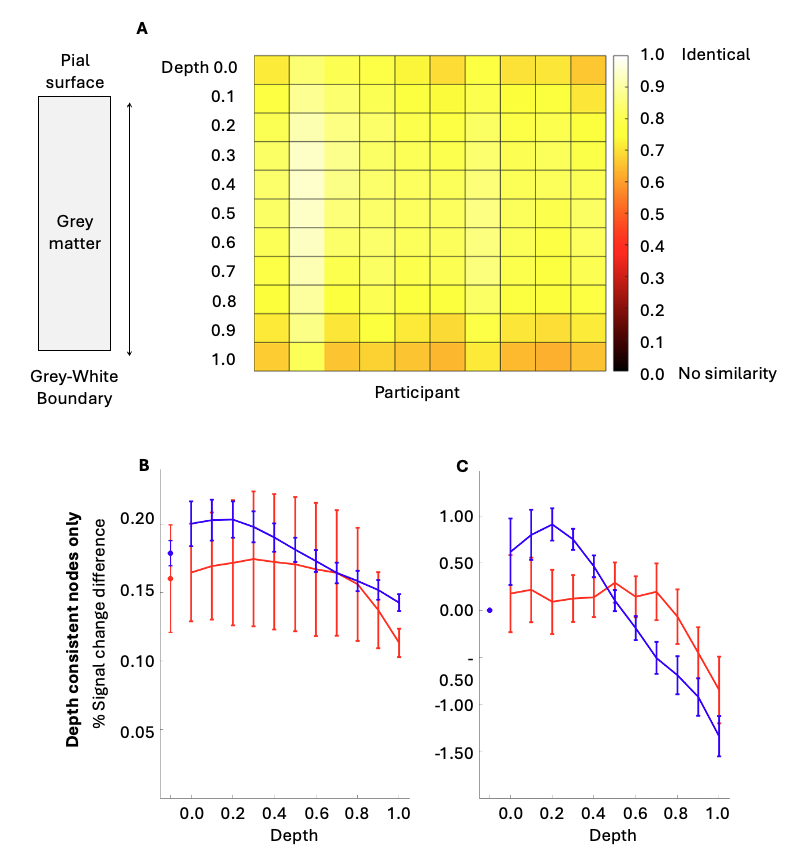

***Fig Supp 6.*** *Additional depth-related figures.* ***A****.* *Comparison of depth-averaged preference maps (Part 1 data) and the preference maps generated at each of the 11 cortical depths (Part 2 data). Each column from left to right represents data from one participant. Each row represents the similarity of that participant’s depth-averaged preference map to each of their 11 depth preference maps. Similarity is calculated using a Jaccard Index, thus a value of 1 indicates complete similarity (i.e., the same 3 Hz/30 Hz stimulation preference at every node) between the depth-averaged preference map and each of the 11 individual depth preference maps. High similarity is indicated by white/yellow colours (see colour bar at the far right).* ***B & C****. Differential signal change over depth for completely depth consistent nodes only. Raw data are presented in the left panel, and z-transformed data on the right panel (to equate means and standard deviations of the distributions, and allow easier visual comparison of shape). This figure is identical to Fig 5 in the main text, but it uses depth-consistent nodes only, where Fig 5 uses all data.*

The depth-averaged preference maps were compared with the 11 individual depth preference maps to determine whether it was appropriate to classify nodes as 3 Hz or 30 Hz preferring using the depth-averaged data when carrying out our laminar analyses. While containing mostly the same data, the laminar data includes the top/bottom 10% of data that was removed from the depth-averaged data. These top and bottom depths were removed from the depth-averaged analyses to reduce the effects of spatial smoothing due to draining veins at the surface of the cortex and the region of lowest signal, respectively. They were included in the laminar analyses to investigate these biases (see *Discussion* for further discussion). As can be seen in *Supp Fig 6* above, mostly light yellow colours are seen, indicating high Jaccard Indices. This suggests that the depth-averaged preference maps are representative of the preference maps over depths, supporting the use of depth-averaged data to split the data into 3 Hz > 30 Hz preferring and 30 Hz > 3 Hz nodes for the laminar analyses.

Comparing *Fig 5* in the main text to *Supp Fig 6* above shows that when only depth-consistent nodes are considered, there appears to be less of a difference in differential signal between 3 Hz preferring nodes and 30 Hz preferring nodes.

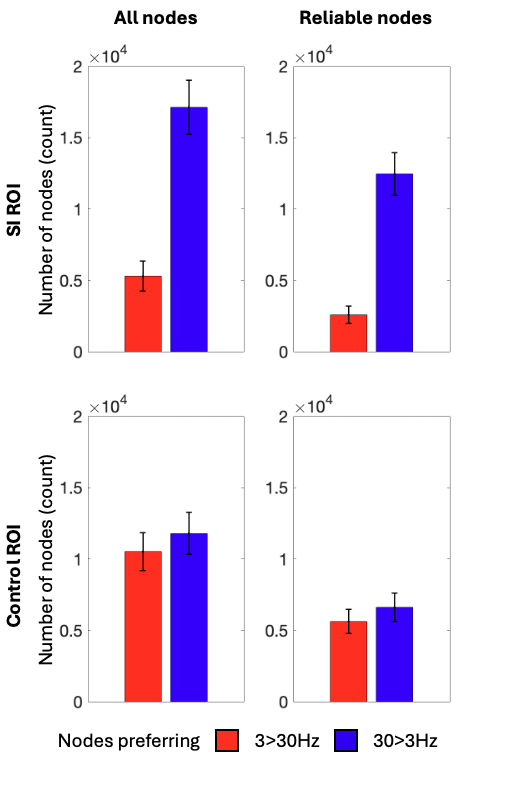

***Fig Supp 7.*** *Difference in preference for 3 Hz versus 30 Hz vibration in the S1 (top panels) and control ROIs (bottom panels), with all nodes (left panels) and just reliable nodes (right panels).*

As can be seen in *Fig Supp 7*, removing unreliable nodes led to a similar reduction in the total nodes preferring 3 Hz and 30 Hz, i.e., there appeared to be a similar proportion of unreliable nodes for both stimulation preferences as before removing unreliable nodes. This was consistently seen in the SI and control ROIs.
